# The investigation on metabolites, genes and open chromatins involved in colored leaves of *Eucommia ulmoides* ‘Ziye’

**DOI:** 10.1101/2022.10.04.510861

**Authors:** Li Long, Shi Qianqian, Yao Wenjing

## Abstract

*Eucommia ulmoides* Oliver ‘Ziye’ has unique purple-red leaves, which contain a variety of flavonoids, so it has high ornamental and medicinal value. However, the categories of flavonoids and molecular mechanism of specific accumulation of flavonoids in ‘Ziye’ leaves is still unclear. Here, differences in metabolic level, gene expression level, chromatin accessibility and cis-regulatory elements were compared between ‘Ziye’ and ‘Huazhong 1’ with green leaf color by metabolome profiling, RNA-seq, and ATAC-seq. A total of 205 flavonoids were identified from these two varieties using ultraperformance liquid chromatography–mass spectrometry (UPLC-MS). The accumulation of most delphinidin, cyaniding, quercetin, myricetin, and isorhamnetin derivatives peaked in old leaves of ‘Ziye’. Single-molecule long-read sequencing indicated that genes in the phenylpropanoid biosynthesis and flavonoid biosynthetic pathway, as well as many transcription factors including MYB, ERF, and WRKY were highly expressed in ‘Ziye’ leaves. ATAC-seq revealed the presence of cell preferentially enriched peaks, which annotated to 6114 genes. Analysis of the genomic regions preferentially accessible in each cell type identified hundreds of overrepresented TF-binding motifs, highlighting sets of TFs such as MYB, ERF, and WRKY that are probably important for color formation of ‘Ziye’ cell. Interestingly, the TFs within each of these cell type-enriched sets also showed evidence of extensively co-regulating each other. Our work demonstrated how chromatin accessibility and TF expression level influenced the expression of flavonoid biosynthesis associated genes, resulted in flavonoids accumulation in ‘Ziye’ leaf. Our results could lay a foundation for further studies of gene expression and functional genomics in *E. ulmoides*.

---

*Eucommia ulmoides* Oliver, the only species in the Eucommiaceae family, is a tertiary relic perennial tree with high economic, ecological and social values because of its ability to produce not only wood but also valuable raw biomaterials for extracting rubber (Wuyun et al., 2017). The leaves and barks of *E. ulmoides* Oliv. are excellent source of flavonoids, chlorogenic acid, lignans, iridoids-α-Linolenic acid and other medicinal ingredients, and have the effects of anti-fatigue, anti-aging, anti-tumor, and enhancing immunity so on. Because the leaves contain a large number of antibacterial and anti-inflammatory active ingredients, it has great potential development in reducing the application of antibiotics in animal feeding. The efficacy of *E. ulmoides* has been well documented in ancient Chinese medicinal books such as Compendium of Materia Medica and Shennong’s Classic of Materia Medica.

*E. ulmoides* ‘Ziye’ is a variety with excellent purplish red leaf. It is bred on the basis of the natural variation of *E. ulmoides* with purplish red leaf. Its purple-red leaf color is so stable that it has the potential to further develop new varieties with multi character aggregation. Many studies suggested that much higher flavonoids especially anthocyanins accumulation mainly contributed to purplish red corloration in *E. ulmoides* ‘Ziye’ leaf, which resulted in much higher medicinal and ornamental value. However, the components and contents of secondary metabolites, and the molecular mechanism of the gradual purple-red color formation of *E. ulmoides* ‘Ziye’ leaf remains to be further studied.

It has been found that the flower and leaf colors are mainly determined by the spatially and temporally restricted deposition of various flavonoids and carotenoids in model plants (Brockington et al., 2015). Flavonoids are widely distributed in plants, and can be divided into seven subclasses including flavones, flavonols, flavandiols, anthocyanins, condensed tannins, chalcones, and aurones. Among these, anthocyanins have the greatest influence on plant tissue pigmentation. The flavonoid biosynthetic pathway can be divided into three steps (Xu et al., 2015; Zhao and Tao, 2015): Firstly, the conversion of phenylalanine to coumaroyl-CoA under phenylalanine ammonia lyase (PAL), cinnamate-4-hydroxylase (C4H), and 4-coumarate CoA ligase (4CL); Secondly, the synthesis of dihydroflavonols from one molecule of coumaroyl-CoA and three molecules of malonyl-CoA catalyzed by chalcone synthase (CHS), chalcone isomerase (CHI), flavanone 3-hydroxylase (F3H), flavonoid 3’5’ -hydroxylase (F3’5’H), flavonoid 3’ -hydroxylase (F3’H), flavone synthase (FNS), and Flavonols synthase (FLS); Thirdly, the synthesis of leucoanthocyanidins from dihydroflavonols under the catalysis of dihydroflavonols 4-reductase (DFR) and then the synthesis of the corresponding colored anthocyanidins by anthocyanidin synthase (ANS) (Xu et al., 2015; Zhao and Tao, 2015). Subsequently, various stable anthocyanins are formed under the catalysis of flavonoid glucosyltransferasen (UFGT) and anthocyanins O-methyltransferase (AOMT) (Lou et al., 2014). The activity of flavonoid biosynthetic enzymes is primarily regulated by a transcriptional activation complex consisting of R2R3-MYB, bHLH ((basic helix–loop–helix) and WD40 (WD40-repeat containing protein).

Transcriptional and metabolic analysis of leaf coloration via RNA sequencing and metabolome has been successfully applied in many plant species including *Pennisetum setaceum* ‘Rubrum’ (Zhu et al., 2020), *Camellia oleifera* (Chen et al., 2017), and red maple (Xia et al., 2021). Thus far, no RNA-seq and metabolome studies of color variation in leaves of *E. ulmoides* have been reported.

Cis-regulatory elements (CREs) are DNA fragments that influence expression of nearby genes through the transcription factor (TF) binding. Proper spatio-temporal activation and silencing of genes is a crucial regulation mechanism involved in various biological processes (Boulay et al., 2018; Rachel et al., 2021). Previously, some CREs in many plants have been general located via enhancer-trapping and QTL mapping (Clark et al., 2006; Wu et al., 2003), but these techniques are time consuming and laborious, excluding extensive genome-wide identification and characterization of CREs. Recently, advances in next-generation sequencing technology including ATAC-seq (Assay for Transposase-Accessible Chromatin) and DNAse-seq have led to improved technical methods for identifying CREs (Boyle et al., 2008; Buenrostro et al., 2013).ATAC-seq is based on the property of TF-binding regulatory regions to form relatively open chromatin, making them sensitive to enzymatic cleavage. The hyperactive Tn5 transposase to fragment DNA while terminally adding adapters at cleavage sites in accessible chromatin regions, which serve for PCR amplification and paired-end next generation sequencing(Crawford et al., 2004; Buenrostro et al., 2013). The advent of large-scale single-cell sequencing has further highlighted the correlation between ATAC-seq identified regions and gene regulation, confirming the utility of this technique in modern plant transcriptomics studies (Farmer et al., 2021; Xu et al., 2021).

Recent studies of ATAC-seq in plants have identified conserved CREs across distantly related species such as Arabidopsis, rice and maize (Oka et al., 2017; Rodgers-Melnick et al., 2016; Sullivan et al., 2014; Zhang et al., 2012). Besides, this technology were used to identify chromatin changes during root-hair differentiation (Maher *et al*., 2018), and compare accessible chromatin regions between pluripotent stem cells in the central zone of the SAM and fully differentiated leaf mesophyll cells that originate from the stem cells of the SAM (Sijacic et al., 2018). However, this study is still absent in chromatin regions identification in *E. ulmoides* ‘Ziye’.

In the present study, we aim to identify the underlying biochemical and genetic evidence of purplish red leaf color variation in *E. ulmoides* ‘Ziye’. Metabolome profiling was performed to identify categories of metabolite, and compare metabolome level between *E. ulmoides* ‘Ziye’ and *E. ulmoides* ‘Huazhong 1’ with green leaves. Single-molecule sequencing data (ONT RNA-seq) were used to explore potential genes involved in metabolite biosynthesis and reveal the molecular mechanisms influencing the development of *E. ulmoides* ‘Ziye’. Furthermore, potential transcription factors, cis-regulatory elements and chromatin accessible regions (ACRs) that may specifically regulate ‘Ziye’ gene expression were identified by ATAC-seq. Our results suggest that distinct classes of TFs collaborate to produce ‘Ziye’ cell type-specific transcriptomes, resulting in flavonoids accumulation especially anthocyanins accumulation in ‘Ziye’ leaf.

## Results

### Anatomical and physiological analyses of ‘Ziye’ leaves

In order to understand the factor that caused the leaf color formation of ‘Ziye’, we measured the content of various colored substances. We firstly determined the contents of anthocyanins and total flavonoids in leaf buds, young leaves, adult leaves, and old leaves. We found that both anthocyanins level and flavonoid level increased gradually with leaf development of ‘Ziye’ and ‘Huazhong 1’, and both of them were much higher in ‘Ziye’ leaves (Figure 1A and 1B). Based on dynamic changes of anthocyanins level and flavonoid level, we selected young leaves (S3 marked in Figure 1A) and old leaves (S6 marked in Figure 1A) which represented for early pigmentation stage and late pigmentation stage for further metabolome, transcriptome and ATAC-seq analysis. P1 and P2 represented for early pigmentation stage and late pigmentation stage of ‘Ziye’ leaves, meanwhile G1 and G2 of ‘Huazhong 1’ leaves were corresponding to the P1 and P2, respectively (Figure 2A).

**Figure 1.**
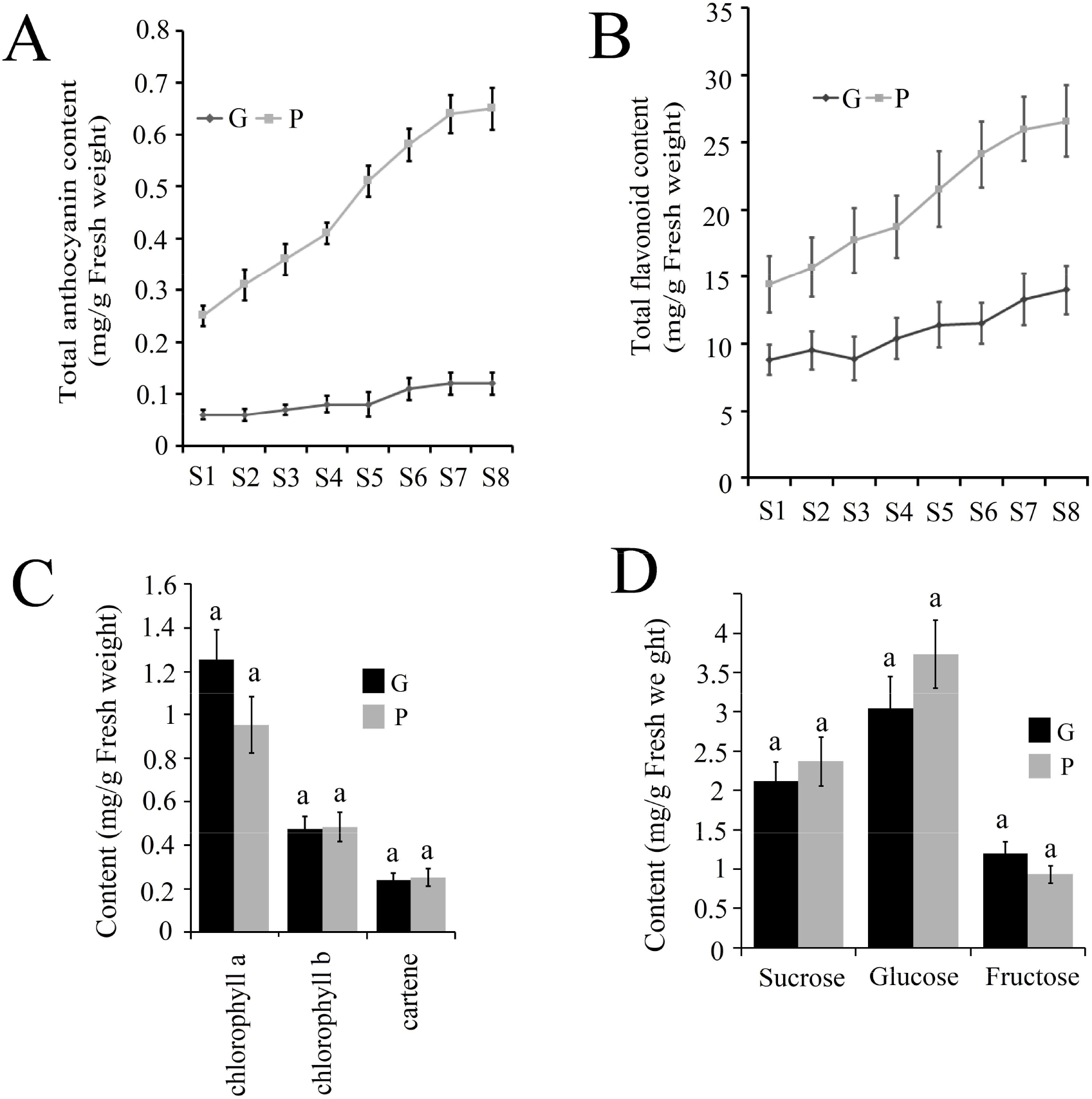
Comparison of pigment contents between ‘Ziye’ *E. ulmoides* and ‘Huazhong 1’*E. ulmoides*. The ratio of (A) total anthocyanins, and (B) total flavonoids to fresh weight during leaf growth and development. The ratio of (C) Chlorophyll and carotene, and (D) soluble sugar to fresh weight in old leaves of ‘Ziye’ and ‘Huazhong 1’.

**Figure 2.**
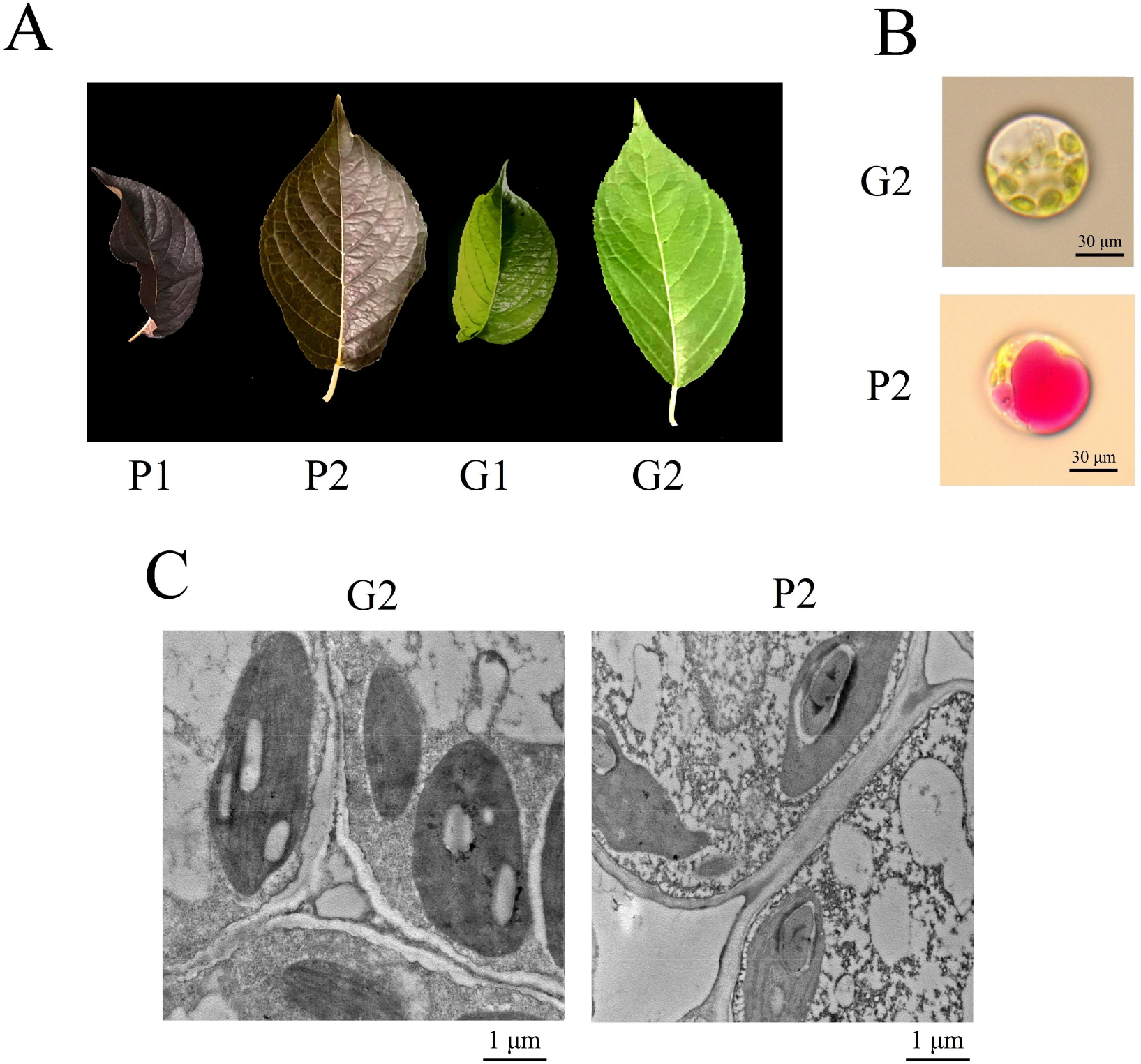
Cellular features of the leaf materials. (A) New fully expanded leaves and old leaves were used for metabolome profiling, isoform sequencing and ATAC-sequencing. (B) Protoplasts from the leaves of the ‘Ziye’ and ‘Huazhong 1’, (C) Cell ultrastructure of the leaves of ‘Ziye’ and ‘Huazhong 1’. G and P represented for *E. ulmoides* ‘Ziye’ and *E. ulmoides* ‘Huazhong 1’, respectively.

Besides, we compared the contents of chlorophyll, carotene, and soluble sugar between the two varieties. The contents of chlorophyll a and fructose were much higher in ‘Huazhong 1’(G2) than those in ‘Ziye’ (P2), but the difference was not statistically significant (Figure 1C and 1D). In contrast, the contents of sucrose and glucose were non-significantly higher in P2 than those in G2. Besides, the chlorophyll b and carotene in P2 were similar to G2.

The leaf color of ‘Ziye’ arised from cytoplasm, not the plastid, and the ‘Ziye’ cytoplasm appeared purplish red under the microscope (Figure 2B). The chloroplast of ‘Ziye’ was larger than that of ‘Huazhong 1’ under the TEM (Figure 2C). In short, anthocyanins and flavonoids enriching in the cytoplasm were the major factor for purplish red color formation in leaves of ‘Ziye’.

### Pigment variation between ‘Ziye’ and ‘Huazhong 1’ leaves

In order to further clarify which components of anthocyanins and flavonoids were the decisive factors for ‘Ziye’ leaf color formation, metabolome profile was carried out using UPLC-MS (Table S1). A total of 205 flavonoids were identified from four *E. ulmoides* leaf samples, including 68 quercetin derivatives, 29 kaempferol derivatives, 2 myricetin derivatives, 4 cyanidin derivatives, 2 peonidin derivatives, 17 delphinidin derivatives, 2 petunidin derivatives. The number of DEMs upregulated and downregulated in P1 vs G1, P2 vs G2 was 104 and 36, 107 and 27, respectively (Figure 3A).

**Figure 3.**
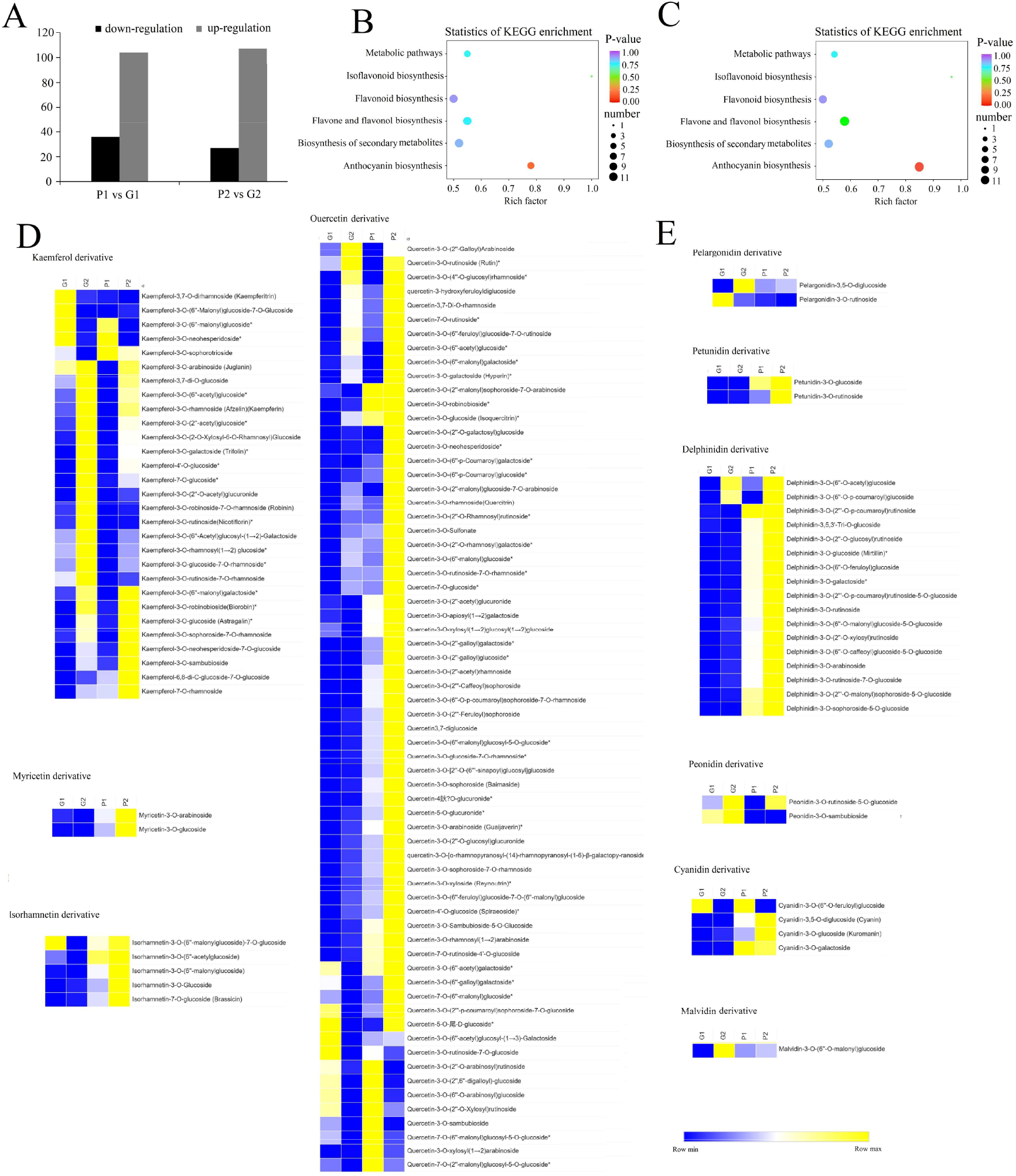
Characterization of metabolite related to ‘Ziye’ leaf color formation. (A) Number of differentially accumulated metabolites. The amount of the differentially accumulated metabolites between P1 and G1, P2 and G2. (B) The KEGG enrichment map of differentially expressed metabolites between P1 and G1. (C) The KEGG enrichment map of differentially expressed metabolites between P2 and G2. (D) Hierarchical clustering heat map of detected flavonols. (E) Hierarchical clustering heat map of detected anthocyanins.

KEGG enrichment analysis indicated that a large amount of anthocyanins were identified as DEMs, of which, most were highly accumulated in P2, indicating the importance of anthocyanins involved in purplish red color formation of ‘Ziye’ (Figure 3B and 3C). Among 27 anthocyanins identified in the leaves of two *E. ulmoides* cultivars (Figure 3E), all two petunidin derivatives and overwhelming majority delphinidin- and cyanidin derivatives were highly accumulated in P2 of ‘Ziye’. In term of peonidin derivatives, ‘Huazhong 1’ leaves mainly accumulated peonidin-3-O-sambubioside, while petunidin-3-O-rutinoside-5-O-glucoside showed similar accumulation level in two cultivators. Two pelargonidin derivatives were highly accumulated in ‘Huazhong 1’. Most flavonols were more abundantly accumulated in ‘Ziye’, especially in P2, except most kaempferols which were highly accumulated in G2 (Figure 3D).

### Overview of single-molecule long-read sequencing results

To identify the key genes involved in ‘Ziye’ leaf color formation, we performed ONT sequencing of ‘Ziye’ and ‘Huazhong 1’ *E. ulmoides* leaves at two different growth stages using the PromethION platform. The number of full-length sequences through primer sequence from two ends of reads obtained from each sample vary from 2,161,952 to 2,951,797 (Table S2). After polishing full-length sequences, redundant-remove analysis through minimap2 software, 68,023 transcript sequences were obtained. These isoforms were grouped into three categories: (1) 26,719 known isoforms that mapped to the known gene set, (2) 31,592 new isoforms of annotated genes, and (3) 9,712 novel isoforms belonging to 4,902 likely new candidate genes that were previously unannotated (Table S3).

### Identification of key pathways involved in ‘Ziye’ leaf color formation

To explore the potential factors that caused the differential coloration and compound content between ‘Ziye’ and ‘Huazhong 1’ leaves, the KEGG enrichment was conducted. The q-value (corrected p-value) of the richness factor was used to evaluate the importance of the pathways involved in purplish red leaf formation (Figure 4).

**Figure 4.**
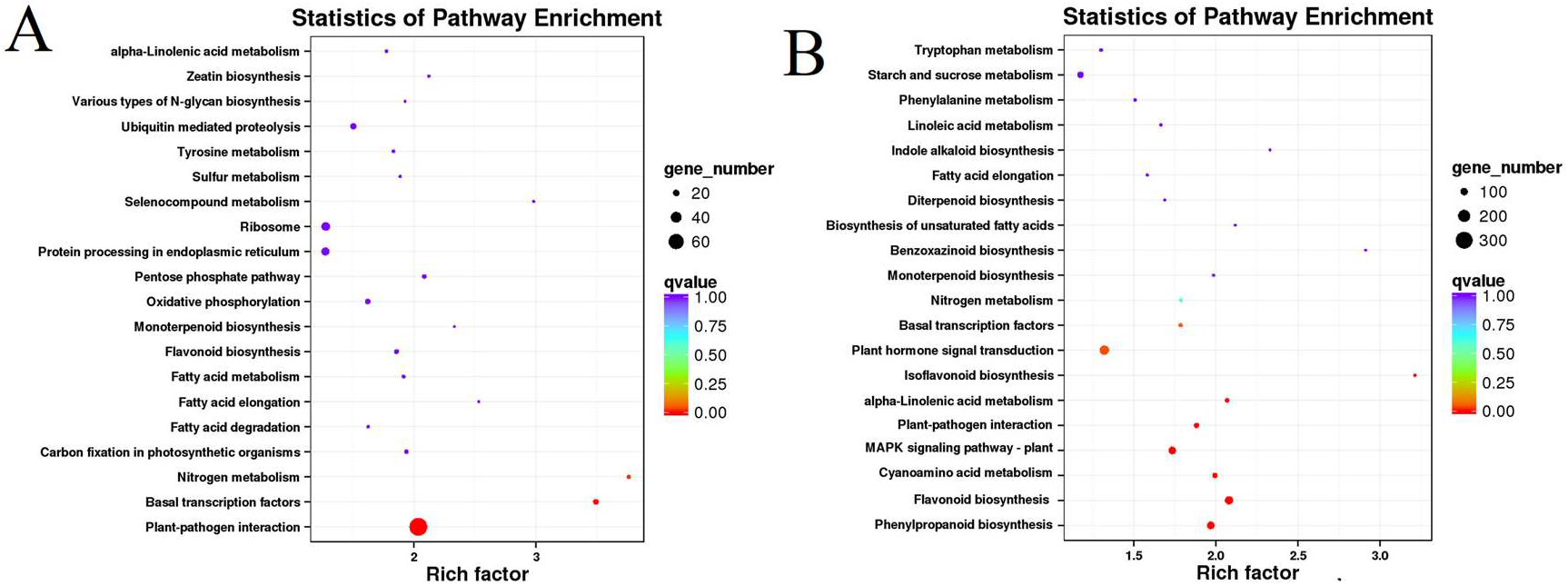
Statistics of pathway enrichment. (A) P1 vs G1, (B) P2 vs G2. The rich factor is the ratio of differentially expressed genes numbers annotated in a given pathway term to all gene numbers that were annotated in the pathway term. The greater rich factor values indicate greater intensiveness.

For G1 vs P1, most “plant-pathogen interaction”, “nitrogen metabolism”, and “basal transcription factor” associated genes differentially expressed (Figure 4A and Table S3). By contrast, the genes involved in “phenylpropanoid biosynthesis” —the precursor for the flavonoids synthesis, “flavonoid biosynthesis”, and “cyanamino acid metabolism” were the most significantly enriched, indicating P2 is the key period for the synthesis of flavonoids in ‘Ziye’ (Figure 4B and Table S3).

### Expression analysis of pigmentation-related flavonoid biosynthetic genes

Based on the annotation results, a total of 43 differentially expressed functional genes were associated with pigmentation-related flavonoid biosynthesis (Table S4). PAL is the first key enzyme of the flavonoid biosynthesis pathway, four of six differentially expressed *PAL* members showed the highest expression level in P2. More than half *4CLs* were highly expressed in the early stage of flavonoids accumulation in ‘Ziye’ (P1). The expression level of *C4M* in both G1 and P1 was higher than that in the P2 and G2. Most CHS members showed high accumulation level in first pigmentation stage of ‘Ziye’ leaves (P1) (Figure 5). One *CHI* (*EUC01961-RA*) who belonged to types I proteins that having CHI enzymatic activity (Morita et al., 2014), showed significantly increasing expression in P2. The *F3′H, FLS* and *F3′5′H* genes played key roles in different flavonols and anthocyanins biosynthetic branches that could underlie various colors. Two differentially expressed *FLS* were identified in *E. ulmoides*. One of them was highly expressed in P1 and the other was highly expressed in P2. Both two differentially expressed *F3’H* were differentially expressed between two color formation stages, but there is no significant difference between the two leaf colors. Among *FLS, EUC20656-RA* showed high expression abundance in both G2 and P2, while *EUC17674-RA* was highly accumulated in P1. *ONT*.*17762*.*1* (*F3′5′H*) showed a high accumulation level in P2.

**Figure 5.**
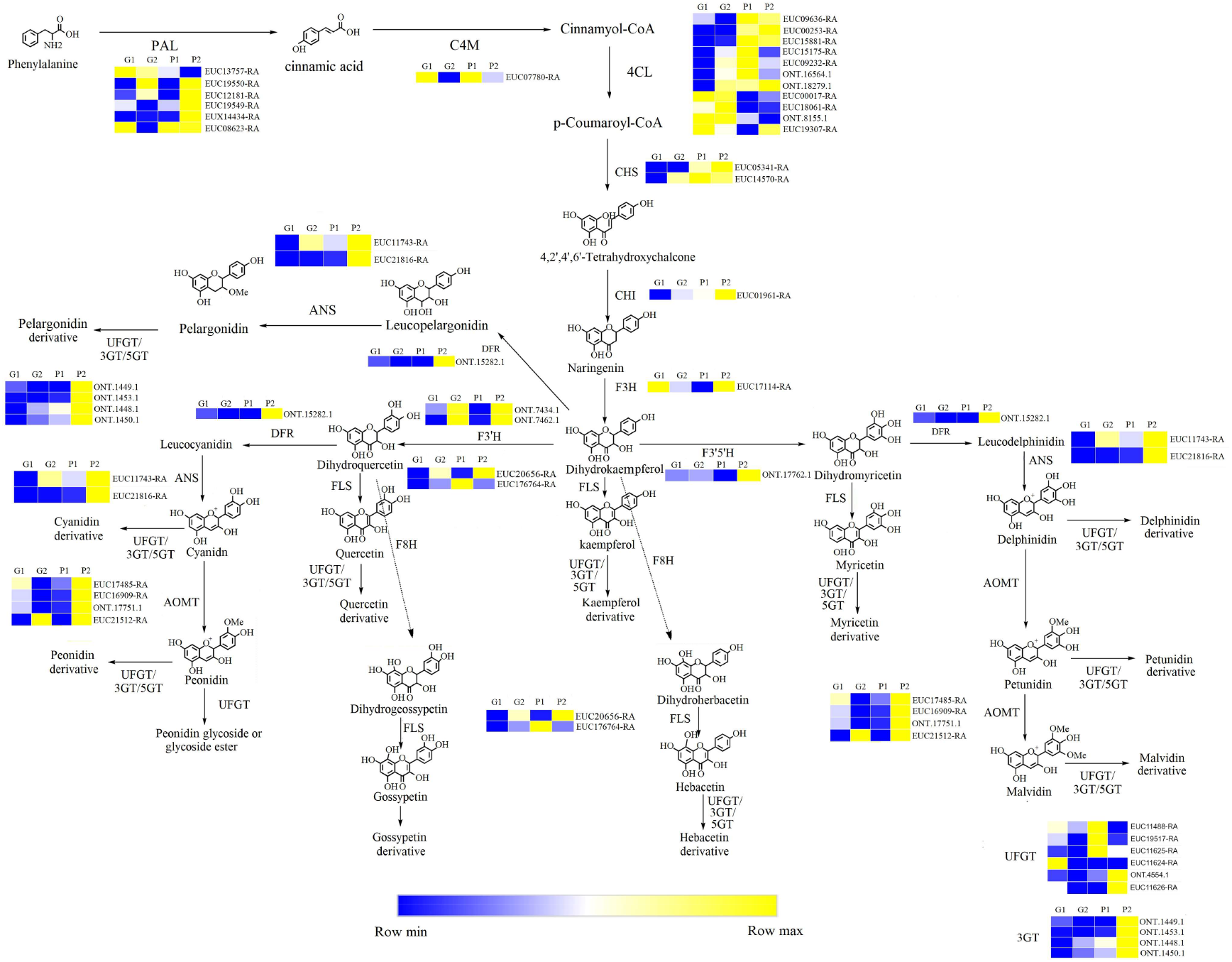
A detailed schematic of the flavonols and anthocyanins biosynthetic pathway and their genes expression levels related to purplish red leaf pigmentation in the *E. ulmoides* ‘Ziye’. Enzyme names and expression patterns are listed at each step. The expression level of each gene in each sample was averaged from three biological replicates of isoform sequencing. Only differentially expressed genes were listed in diagram. White indicates moderate expression level, blue indicates low accumulation level, and yellow indicates high accumulation level.

There were two main branches from dihydrokaempferol into different anthocyanins. Cyanidin and delphinidin formed from dihydroquercetin and dihydromyricetin and catalyzed by DFR and ANS, respectively. Here, one *DFR* and two *ANS* genes were differentially expressed, and all these three genes were highly expressed in P1 or P2. Later, peonidin-based anthocyanins were converted from cyanidin through a series of glycosylation and methylation reactions to produce colors. In another anthocyanins biosynthetic pathway, malvidin and petunidin were converted from delphinidin through a series of glycosylation and methylation reactions. The methylation and glycosylation reactions in both two anthocyanins synthesis branches mainly regulated by anthocyanins O-methyltransferase (AOMT). Four *AOMTs* differentially expressed during ‘Ziye’ leaf pigmentation, and all of them performed high accumulation level in P2. Most *UFGT* genes, which played key roles in 3-glucoside formation in different flavonols and anthocyanins biosynthetic branches, showed higher expression levels in P1 and P2.

### The expression changes of transcription factor in the ‘Ziye’ pigmentation process

Nextly, we focused on differentially expressed transcription Factors (TFs) (Figure 6). More than half differentially expressed ERF, WRKY, NAC were highly accumulated in P2, suggesting that these TFs could play important roles in leaf pigmentation. Among 155 genes annotated as R2R3-MYB TFs, 20 genes were identified as DEGs. More than one third of the differentially expressed R2R3-MYB expressed almost exclusively in P2. Of these, *EUC08247-RA* (*MOF1*), *EUC03590-RA* (*MYB78*), and *EUC10567-RA* (*MYB30*) were mostly up-regulated. Seven *MADS*, five *bZIP*, and one *WD40* differentially expressed during ‘Ziye’ leaf pigmentation. *EUC03326-RA* (*AGL8*), *EUC06393-R*A (*EJ2*), *EUC09890-RA* (*bZIP53*), *EUC01992-RA* (*bZIP16*) and *EUC22218-RA* (*HOS15*), which showed high accumulation level in P2, P2, P1, P1, and P1, respectively, might be associated with purplish red leaf formation.

**Figure 6.**
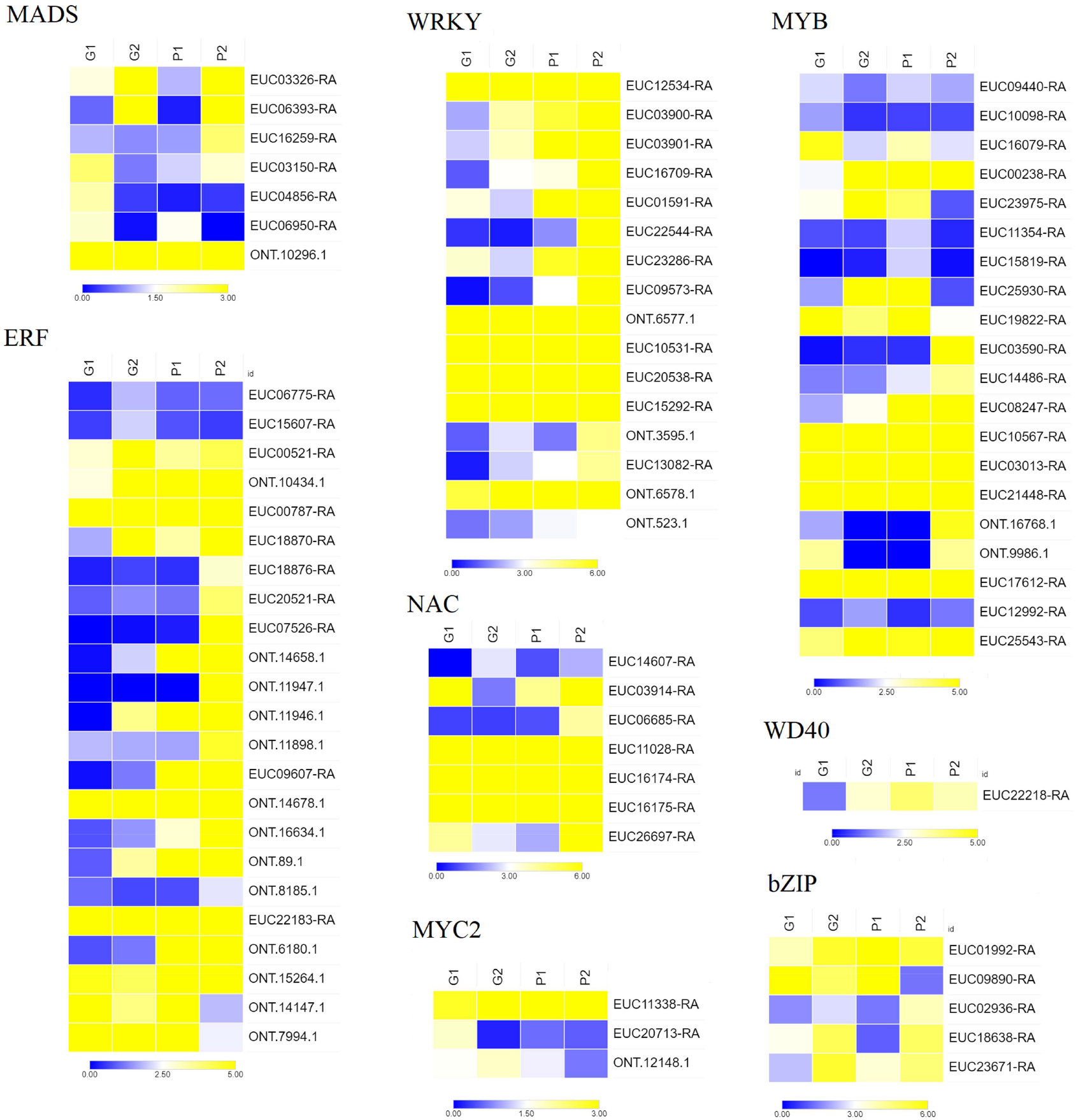
Expression profiles of differentially expressed transcription factors involved in purplish red leaf pigmentation in the *E. ulmoides* ‘Ziye’. Only differentially expressed genes were listed in diagram. White indicates moderate expression level, blue indicates low accumulation level, and yellow indicates high accumulation level.

### Identification of ‘Ziye’ leaf-specific accessible chromatin regions and TF binding motifs

To examine the local chromatin landscape of ‘Ziye’ leaves, and compare the difference between the ‘Ziye’ leaves and ‘Huazhong 1’leaves in the open chromatin region, we performed ATAC-seq. More than 86 million reads were obtained for each biological replicate through paired-end sequencing (Table S5 and S6). After aligning the ATAC-seq reads to the *E. ulmoides* genome, we found that, on average, 30.66% of all reads were uniquely mapped to the genome. The accessible regions identified were mostly enriched around the TSS (Figure 7A) in both two leaves, which was consistent with these regions containing cis-regulatory elements in *E. ulmoides*, and also consistent with previous reports in other plants (Rachel et al., 2021; Dai et al., 2022). The genomic distribution of reproducible THSs (Transposase hypersensitive sites) is similar between ‘Huazhong 1’and ‘Ziye’, with 41.47% (G2) and 40.12% (P2) of THSs located within 1 kb upstream of the gene transcription start site (TSSs), 19.44% and 17.22% located within 1 kb – 2 kb upstream of the TSSs, 8.04% and 9.04% located within 2 kb – 3 kb upstream of the TSSs, 10.09% (G2) and 10.02% (P2) located within the gene body, 40.96% (G2) and 43.20% (P2) located in the intergenic region (Figure 7B). This genomic distribution of reproducible THSs suggested that the majority of cis-regulatory regions in the *E. ulmoides* genome were located in the vicinity of gene core promoters, as previously observed in Arabidopsis (Lu et al., 2016).

**Figure 7.**
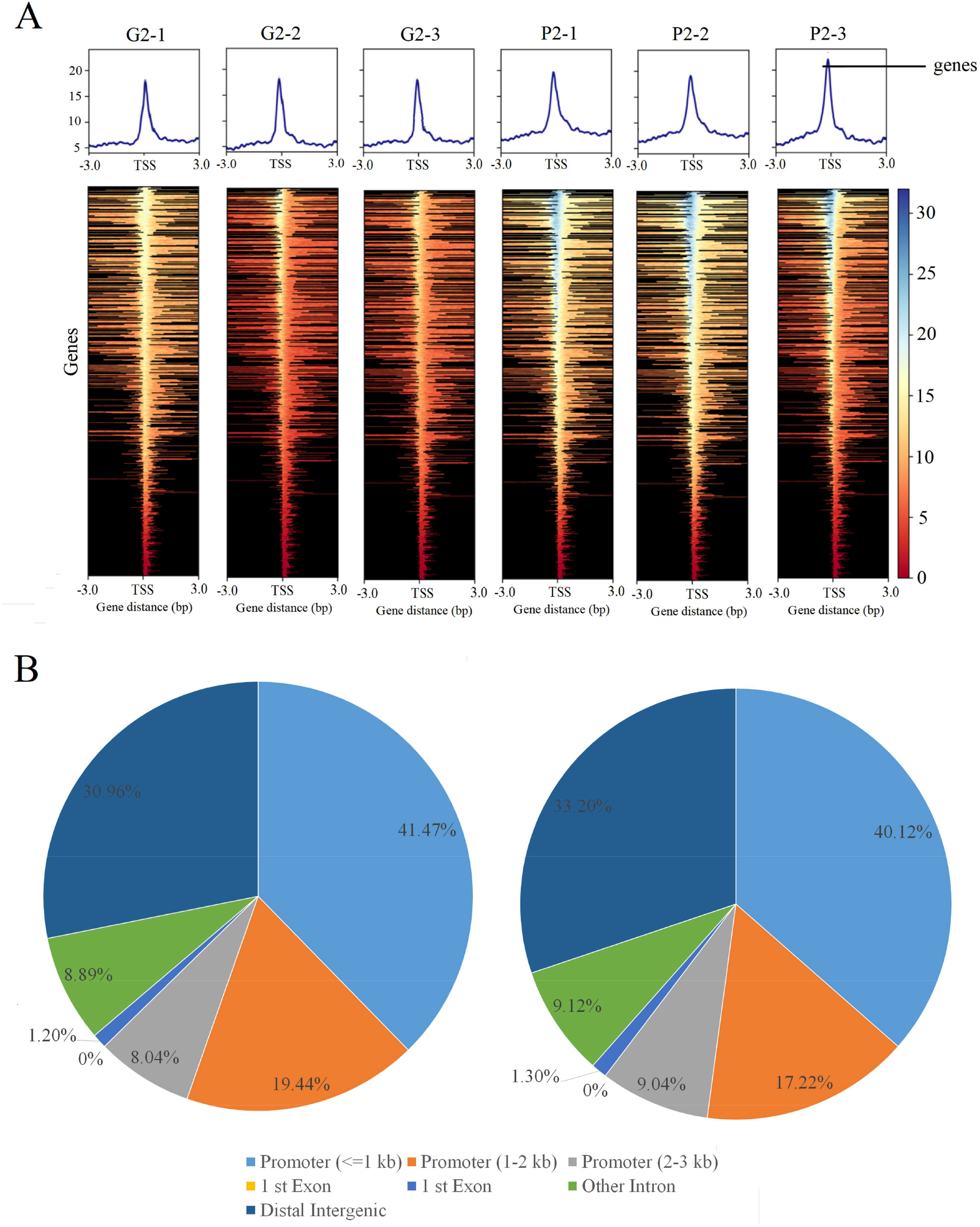
Distribution of accessible regions were identified using ATAC-seq. (A) Distribution of accessible regions around the TSS (Transcription start sites) identified from ATAC-seq using the leaves of *E. ulmoides* ‘Huazhong 1’ and *E. ulmoides* ‘Ziye’. (B) A pie chart showed the distribution of accessible regions identified using ATAC-seq from the sample of ‘Huazhong 1’and ‘Ziye’ leaf throughout the *E. ulmoides* genome.

To determine the regulatory landscape of the *E. ulmoides* genome between ‘Ziye’ and ‘Huazhong 1’cells, we generated genome-wide cell type-preferentially enriched maps of accessible chromatin regions (ACRs) from ATAC-seq. In total, 289,300 peaks and 244,953 peaks were identified in the ‘Ziye’ and ‘Huazhong 1’ cells, respectively. Among these, 13770 cell preferentially enriched peaks (differentially accessible chromatin regions, dACRs) were found and were annotated to 6114 genes (Table S7). 30.3 % of DEGs identified by transcriptome sequencing had differential ACRs between ‘Ziye’ and ‘Huazhong 1’cells. These genes were enriched in the biological process of response to light stimulus, abscisic acid-activated signaling pathway, proteasome-mediated ubiquitin-dependent protein catabolic process, response to stress. These results suggested that opening or closing of nuclear chromatin region of genes related external environmental stimulation especially illumination were essential for the color formation of ‘Ziye’ leaf (Figure 8A).

**Figure 8.**
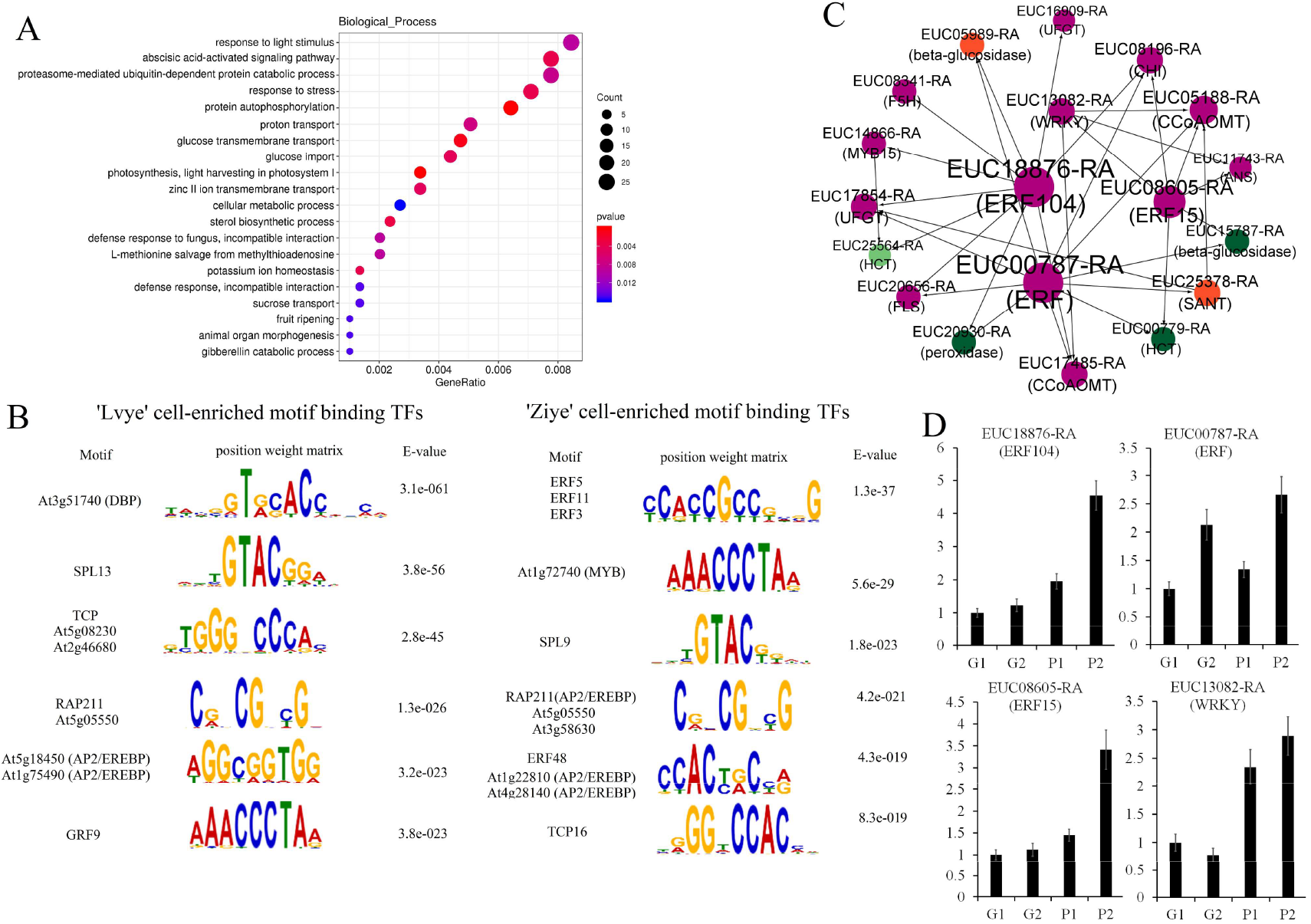
(A) The KEGG enrichment map of differential peak region related genes. (B) The six most enriched position weight matrix (PWM) of transcription factor binding sites, and E-value from the MEME-ChIP analysis were shown for ‘Huazhong 1’cells (left) and ‘Ziye’ cells (right). (C) The putative regulatory networks for three ERF genes highly expressed in ‘Ziye’ were constructed. Each TF circle has regulatory inputs (TF-binding site within its proximal regulatory regions) and regulatory outputs (that TF’s predicted binding site in the other gene’s proximal regulatory regions). For example, EUC13082-RA (WRKY) has Two regulatory outputs to EUC17485-RA (CCoAOMT), and EUC05188-RA (CCoAOMT), and one regulatory input to EUC08605-RA (ERF15). Light green, dark green, orange and purplish red represent the isoforms with the highest expression levels in young leaves of ‘Huazhong 1’ (G1), old leaves of ‘Huazhong 1’ (G2), young leaves of ‘Ziye’ (P1) and old leaves of ‘Ziye’ (P2), respectively. Larger edges in the network indicated geneedges with more connections.

From ACRs identified by our ATAC-seq, MEME-ChIP discovered 131 centrally enriched motifs which might be recongized and binded by TFs (Figure 8B). A large amount of these motifs related to ERF (ERF5, ERF11, ERF3, RAP211, ERF48 et al.) family, MYB (MYB58, MYB88, LHY1, et al) family, and WRY family, bZIP (bZIP18, bZIP42, bZIP43, et al) et al, were enriched in ‘Ziye’. For ERF as an example, a total of 87 *ERFs* expressed in the *E. ulmoides* leaves and 23 showed differential expression between ‘Ziye’ and ‘Huazhong 1’ cells (Figure 6). Interestingly, nearly three-quarter of 29 differentially expressed *ERFs* were preferentially expressed in the ‘Ziye’ cells, which may explain the overrepresentation of the ERF binding motif in ‘Ziye’ cells. Their binding sites widely distributed in promoters of flavonoid biosynthesis related genes, such as EUC18876-RA (ERF104) which showed highest accumulation level in P2, targeted to *EUC08196-RA* (*CHI*), *EUC20656-RA* (*FLS*), *EUC17485-RA* (*AOMT*), *EUC14866-RA* (*MYB15*) as well as many phenylpropaniod biosynthesis genes (Figure 8C). Most of these target genes performed high accumulation level in P2, indicating that ERFs played critical roles in regulating purplish red leaf pigmentation. Similar results were found in MYB, WRKY, bZIP families. Other common TF binding sites in ‘Ziye’ cells included those for C2H2, basic helix-loop-helix (bHLH34, bHLH104, et al.), and others. The significant enriched TF binding sites in ‘Huazhong 1’cells related to DBP, SPL13, TCP, et al.

## Discussion

### Flavonols and anthocyanins accumulations affect color in ‘Ziye’ leaves

The determination of pigment content indicated that the formation of cytoplasmic color is mainly attributed to higher accumulation level of flavonols and anthocyanin. In the process of plant color formation, malvidin is responsible for blue-colored tissues, cyanidin is abundant in reddish-purple (magenta), delphinidin appears blue-reddish and purple colors, petunidin has been detected in purple tissues, pelargonidin for red color, and peonidin for magenta color (Robinson and Robinson, 1932; Tanaka and Ohmiya, 2008). These six types of anthocyanins were all detected in ‘Ziye’ leaves. Among these, cyaniding, petunidin and delphinidin derivatives were predominant in ‘Ziye’ leaves, indicating that these anthocyanins may be responsible for the purplish red leaves. Besides, quercetin, isorhamnetin, myricetin which were closely related to the color appearance in petals of lily cultivars and *Primula vulgaris* cultivars (Lei et al., 2017, Wu et al., 2016, Li et al., 2020) were also highly accumulated in ‘Ziye’. These findings differed from some previous studies. For example, cyanidin, pelargonidin, delphinidin, and peonidin had important functions in color formation of *Camellia japonica* petals (Fu et al., 2021); cyanidin, pelargonidin, and delphinidin were the main anthocyanins in the red flower petals of strawberries (Xue et al., 2019); malvidin contributed mostly to the red petals of *Bauhinia variegate* (Zhang et al., 2021). Whereas in our study, malvidin and peonidin derivatives—the two common types of anthocyanins, which caused plant pigmentation, were not the main reason for the purplish red formation of *E*.*ulmoides* leaves. In fact, delphinidin, petunidin, cyaniding, quercetin, myricetin, and isorhamnetin derivatives were the main pigments that contributed to the purplish red coloration.

### Key structural genes responsible for flavonols and anthocyanins biosynthesis in ‘Ziye’

In the anthocyanins and flavonols biosynthetic pathway, PAL is one of the key enzyme families (Huang et al., 2010). Expression analysis showed that most *PAL*s exhibited high abundances in old leaves of ‘Ziye’, associated with high accumulation of the product of these metabolites in old leaves of ‘Ziye’. CHS and CHI have important effects on the accumulation of flavonoids. A previous study reported that in untransformed tobacco flowers, anthocyanins was not accumulated, while the accumulation of flavonoids increased by over-expression of *CHI* from chrysanthemum (Li et al., 2006). Overexpression of peony *CHI* in tobacco also increased the accumulation of flavonoids (Zhou et al., 2014). Cyanidin and delphinidin derivatives represent two independent branches from dihydrokaempferol, which are catalyzed by *F3’H* and *F3’5’H*, respectively (Li et al., 2020). In cyanidin derivatives biosynthesis branch, the relatively balanced expression of *F3’H* between ‘Ziye’ and ‘Huazhong 1’ leaves resulted in no significant difference in the content of Dihydroquercetin (Taxifolin) between these two leaves. In the next step, cyanidin was abundantly accumulated in old leaves of ‘Ziye’, which was biosynthesized from dihydroquercetin and leucocyanidin catalyzed by DFR (ONT.15282.1) and ANS (EUC11743-RA and EUC21816-RA). Meanwhile, DFR and ANS were also highly expressed in old leaves of ‘Ziye’. In delphinidin derivatives branch, the high accumulation level of delphinidin derivatives were consistent with the expression profiles of *F3’5’H*. On the whole, most differentially expressed *CHS, CHI, F3H, F3’H, ANS, DFR*, and *FLS* genes in current study were positively correlated with differentially accumulated flavonols, anthocyanins as well as metabolic intermediates in the comprehensive analysis of the metabolome and transcriptome, and these genes performed much higher accumulation levels in ‘Ziye’, which is consistent with results in the purple-leaved Jujube and *Hopea hainanensis* (Li et al., 2021; Huang et al., 2022). These results suggested that the high expression of these structural genes could promote the accumulation of flavonols and anthocyanins. The final step in the anthocyanins and flavonoids biosynthesis pathway is 3-glucoside formation by UFGT, which is the key enzyme for anthocyanins and flavonoids stability. The expression of *UFGT* genes were detected in ‘Ziye’ cultivars, but not in ‘Huazhong 1’, which consisted with previous report in Arabidopsis that AtUDT78D2 (flavonoid 3-O-glucosyltransferase) catalyzes the glycosylation of both flavonols and anthocyanins, were associated with anthocyanins and flavonols accumulation in purplish red leaves (Yonekura-Sakakibara et al., 2008).

### The regulation network involved in leaf color formation of ‘Ziye’

In order to find key TF families regulating ‘Ziye’ color formation, we analyzed the differentially enriched chromatin accessible regions to identify putative cell type-specific cis-regulatory motifs as well as the TFs that bind them (Figure 3a). We found that chromatin accessibility, which was positively associated with the expression level of nearby genes in both ‘Ziye’ and ‘Huazhong 1’cells, contributed significantly to the cell type-preferentially enriched expression of DEGs, as 30.3 % of DEGs had differential ACRs between ‘Ziye’ and ‘Huazhong 1’cells, including the key genes involved in flavonoid biosynthesis, phenylpropaniod biosynthesis, and environment stimulation. Thus, the differentially enriched THS between ‘Ziye’ and ‘Huazhong 1’ played important roles in regulating the gene expression during purplish red leaf color formation process.

Using transcriptome sequencing data for these two cell types, we found many TFs that were highly expressed in ‘Ziye’ cell type and whose motifs were also overrepresented in THSs enriched in that cell type such as ERF, MYB, WRKY, bHLH et al. In all species studied to date, flavonoid regulation is mainly controlled by a protein complex formed by R2R3-MYBs, bHLH and WD40, which regulate the expression level of flavonoid biosynthetic genes (Xu et al., 2014; Zhang et al., 2017). In current study, the importance of MYB and bHLH were also confirmed by our RNA-seq and ATAC-seq data. Interesting, we also found that large amount of ERF and WRKY were highly accumulated in ‘Ziye’ and many ERF and WRKY binding motifs were enriched in ACRs of ‘Ziye’ cells. The putative regulatory networks analysis showed three ERFs which showed highest expression level in ‘Ziye’, their binding motifs were widely distributed in open chromatin region related to phenylpropaniod biosynthesis genes and flavonoid biosynthesis genes and many TFs. For example, EUC18876-RA targeted to EUC14866-RA (MYB15), EUC08605-RA targeted to EUC13082-RA (WRKY), and EUC14866-RA and EUC13082-RA simultaneously regulate some other anthocyanin biosynthesis genes expression. These TFs and functional genes formed a complex regulatory network and participated in anthocyanins biosynthesis in ‘Ziye’ leaves.

Following the logic that the identified TF motifs probably represent true TF-binding events when they occur within an open chromatin region of the corresponding cell type, we were able to predict the target genes of TFs of interest (Figure 4a). Based on GO annotation, a large amount of cell preferentially enriched peaks was found in promoters of genes related to light stimulus, abscisic acid-activated signaling, and other external environmental stimulation, partially agree with DEGs identified by transcriptional sequencing. Many studies suggested that light, especially UV-B, was an important environmental factor to promote the accumulation of anthocyanins (Ma et al., 2021; An et al., 2018; Kumar et al., 2013). In red pear, Ethylene-activated PpERF105 inhibit anthocyanins biosynthesis by induces the expression of the *PpMYB140* which act as a repressor in anthocyanins biosynthesis (Ni et al., 2021). The WRKYs involved in flavonoid biosynthesis were also reported in many model plants (Verweij et al., 2016; Hu et al., 2020). In current study, many ERF, WRKY, bZIP, NAC annotated as environmental stimulation by GO, with ‘Ziye’ cell preferentially enriched peaks occurred in their promoter region, such as EUC12534-RA (WRKY) annotated as response to drug (GO:0042493), response to oxygen-containing compound (GO:1901700), and response to chemical (GO:0042221); EUC02329-RA annotated as cellular response to endogenous stimulus (GO:,0071495), and response to external biotic stimulus (GO:0043207), indicating chromatin opening stimulated expression of these TFs. In short, chromatin opening influenced expression of both functional genes and TFs.

In conclusion, many ACRs specially enriched in ‘Ziye’ meaning the DNA sequence of these ACRs would specially bound by many TFs, including ERF, MYB, WRKY, bHLH et al. The high abundance of these TFs in ‘Ziye’ and overrepresented of binding sites related to these TFs in THSs of promoters in ‘Ziye’ stimulated the expression of the flavonols and anthocyanins biosynthesis genes, which eventually leads to the accumulation of delphinidin, cyaniding, petunidin, quercetin, myricetin, and isorhamnetin derivatives in ‘Ziye’ cytoplasm.

## Methods

### Plant material

Grafted seedlings of *E. ulmoides* ‘Ziye’ with purplish red leaves and *E. ulmoides* ‘Huazhong 1’ with green leaves were planted at the nursery of Northwest A&F University, Yangling, Shaanxi. Two-year-old saplings of ‘Ziye’ and ‘Huazhong 1’ were used for sample collection. young leaves of ‘Ziye’ (P1), young leaves of ‘Huazhong 1’ (G1), old leaves (20 days after full expansion) of ‘Ziye’ (P2), and old leaves of ‘Huazhong 1’ (G2) were harvested separately for further transcriptomic, metabolomics, and ATAC-seq analysis. For metabolite profiling, transcriptome sequencing, and ATAC sequencing, five leaves from each individual and each stage were collected from tetrad independent individuals to mixed into one biological replicate. Three biological replicates were prepared for each test.

### Protoplast preparation

The preparation of free protoplasts from palisade tissues were performed according to previously described methods (Yoshida et al., 2009; Qi et al., 2013). The released protoplasts were observed under a photographic microscope (Olympus CX41, Tokyo, Japan).

### Transmission electron microscope

After 3% glutaraldehyde fixation, all leaf samples were washed with 0.1 M PBS for five times, postfixed with 1% osmic acid for 3 h, and sequentially dehydrated using a graded acetone series (30%, 50%, 70%, and 90%). Spurr-embedded tissues were sectioned by an ultramicrotome (LEICA UC6, Weztlar, Germany) and then further observed with a transmission electron microscope (JEM-123O, Tokyo, Japan).

### Sample preparation and metabolomic analysis

Metabolite analysis was performed using an ultra performance liquid chromatography-electrospray ionization-tandem mass spectrometry system (UPLC-ESI-MS/MS). Different leaf samples were rapidly snap-frozen in liquid nitrogen and stored at -80 °C until further processing. Leaf samples were freeze-dried by vacuum freeze-dryer (Scientz-100F, Scientz, Ningbo, China), and crushed using a mixer mill (MM 400, Retsch, Haan, Germany), subsequently. 0.1 g powdery tissue was dissolved with extraction buffer, and then placed in a refrigerator at 4 °C overnight. The extracting solution was filtrated by centrifugation at 12000 rpm for 10 min before UPLC-MS/MS analysis.

The extracting solution was analyzed using an UPLC-ESI-MS/MS system. We selected waters ACQUITY UPLC HSS T3 C18 column (1.8 μm, 2.1 mm*100 mm) as the chromatographic column. Mobile phases A was pure water with 0.1% formic acid, and mobile phases B acetonitrile with 0.1% formic acid, respectively. The column oven was maintained at 40 °C, and the injection volume was 4 μL. Sample measurements were performed with a gradient program that employed the starting conditions of 95% A and 5% B at 10 min, a linear gradient to 5% A and 95% B was kept for 1 min. Subsequently, a composition of 95% A and 5% B was adjusted within 1.1 min and kept for 2.9 min. The flow velocity was set as 0.35 ml per minute. The effluent was alternatively connected to an ESI-triple quadrupole-linear ion trap (QTRAP)-MS.

The detection of metabolites eluted from the column was executed by ESI-triple quadrupole-linear ion trap (QTRAP)-MS (ABI, Waltham, MA, USA), in both positive and negative polarity mode. The ESI source operating parameters, instrument tuning and mass calibration were referenced to previous work (Cho et al., 2016). Based on the optimized collision energy and declustering, the triple quadrupole scan (QQQ) was used to scanning each ion pair (Chen et al., 2013). The absolute metabolite was quantified by ANOVA, and Tukey’s honest significance difference test was used for paired comparison via PAST v.3. x (Hammer et al., 2001).

### RNA extraction and quality assessment

For each sample, total RNA extraction and separation of residual DNA contamination were excuted using Plant RNA Extraction Kit (TaKaRa MiniBEST, kyoto, Japan), and RNase-free DNase I (Thermo Fisher Scientific, UK), respectively Then, 1.5% agarose gels, and an Agilent Bioanalyzer 2100 system (Agilent Technologies, Santa Clara, CA, USA) were carried out in order to check the purity and integrity of the RNA.

### ONT library preparation, sequencing and analysis

1µg total RNA was prepared for cDNA libraries using cDNA-PCR Sequencing Kit (SQK-PCS109, Oxford Nanopore Technologies, Oxford, UK). The final cDNA libraries were added to FLO-MIN109 flow cells and run on PromethION platform (Biomarker Technology Company, Beijing, China). Raw reads generated from the PromethION platform were filtered with the default parameters (average read quality score ≥ 7; read length ≥ 500bp). Ribosomal RNA was removed after mapping to rRNA database.

Next, full-length, non-chemiric (FLNC) transcripts were determined by searching for primer at both ends of reads. Clusters of FLNC transcripts were obtained after mapping to *E. ulmoides* reference genome with Minimap2 (Wuyun et al., 2017), and consensus sequences were obtained after polishing within each cluster by Pinfish. In order to remove redundant, consensus sequences were mapped to *E. ulmoides* reference genome using minimap2. Mapped reads were further collapsed by cDNA_Cupcake package with default parameters (identity ≥ 90%; min-coverage ≥ 85%). Different transcripts caused by different 5’ sequences were not taken into consideration when collapsing redundant transcripts.

Full length reads were mapped to the reference transcriptome sequence. Reads were further used to quantify if the match quality was above 5. Expression levels were calculated by reads per gene/transcript per 10,000 reads mapped (CPM). Differential expression analysis of two experimental groups was performed using the DESeq2 R package (1.6.3) (Anders et al., 2010). Genes with a FDR < 0.01 and fold change ≥ 2 were assigned as differentially expressed.

All the identified genes were annotated using the Kyoto Encyclopedia of Genes Genomes (KEGG) and Gene Ontology (GO) databases with BLAST and a p-value was set as 10^−7^. GO enrichment analysis was conducted using TBtools with Fisher’s exact test (Chen et al., 2018).

### Expression profile analysis by qRT-PCR

Total RNAs were reversed into first-strand cDNA using the M-MLVRT reverse transcription kit (Promega, USA). qRT-PCR was performed with SYBR Premix Ex Taq (TaKaRa Biotech Co., Ltd., Dalian, China) on CFX96 Connect real-time PCR detection system (Bio-Rad Laboratories, Inc., Hercules, CA, USA) according to the manufacturer’s instructions. The relative expression levels of genes were normalized to the *Ubiquitin-conjugating enzyme E2* (*UBC E2*) expression level in the same sample and calibrated to the transcription level in the ‘Huazhong 1’ at G1 (Ye et al., 2018). Each qRT-PCR assay was performed in three independent biological replicates and three technical replicates. All primers were designed by by Oligo 7 software (Table S8).

### ATAC-seq

The old leaves of ‘Ziye’ and ‘Huazhong 1’ were used for ATAC-seq. For each experiment, 50000 cells were centrifuged with 500 g centrifuging parameters for 5 min under 4 °C, and then the supernatant was removed. The cells were washed with 0.1 M PBS, and then suspended with cold lysis buffer. The supernatant was removed using 4 °C, 10 min and 500 g centrifuging parameters. The transposing reaction system was configured with the Tn5 Transposase. The cell nucleus was suspended with the transposing reaction system, and the DNA were purified after incubating at 37 °C for 30 minutes. The PCR reaction system was configured with the purified DNA, and then the PCR amplification was performed. The DNA libraries were sequenced on Illumina platform, finally.

Illumina Sequenced Reads Processing Raw Reads were filtered to remove adapters (the adapters and reads less than 35bp) and low quality reads, and then high-quality Clean Reads provided in FASTQ format were obtained for subsequent analysis. The Bowtie2 software was used to compare the high-quality Reads with the reference genome to obtain the alignment efficiency of the sample Reads and the position information of the Reads on the genome (Langmead et al., 2012). DeepTools v2.07 was used to map the density distribution of sequencing Read in the 3 kb of interval upstream and downstream of the TSS (transcription start site) of each gene, and the results were presented as heat maps (Fidel., 2015). The Reads abundance in the whole genome was statistically analyzed by the sliding window method. FDR < 0.05 was selected as the identified peak via MACS2 v2.1.1 software (Zhang et al., 2008). The ChIPseeker package was used for functional annotation of genome-wide Peak (Yu et al., 2015). The average Peak number of the TSS region was counted. The Peak region was annotated as to whether the Peak region was in the 5 ‘UTR, 3’ UTR, TSS, Intronic, Exon or Intergenic region according to the distance relationship between the Peak region and the functional elements of each gene on the genome. The Peak region associated genes were annotated by NR (Non-Redundant Protein Sequence Database), Swiss-prot, GO, KEGG, COG, KOG, eggNOG and Pfam database. DiffBind package was used for Difference Peak analysis (fold change > 1.5, P-value < 0.05) (Stark et al., 2011). According to the distance relationship between the differential Peak regions and the functional elements of each gene on the genome, the differential Peak region was annotated with the functional databases mentioned above.

The differentially accessible chromatin regions (dACRs) that significant enriched in ‘Huazhong 1’ cell or ‘Ziye’ cell were considered as cell specific type, and were further used for motif analysis. The cell type-enriched Peak region from each cell type were first adjusted to the same size (200 bp). The sequences present in these scaled regions were isolated using the Regulatory Sequence Analysis Tools (RSAT), which also masks any repeat sequences (Medina-Rivera et al., 2015). The masked sequences were run through MEME-ChIP with default parameters to identify motifs that were present in higher proportions than expected by chance (E-value ≤ 0.05) (Machanick and Bailey, 2011).

### Accession numbers

We deposited the RNA-seq data and ATAC-seq in the Sequence Read Archive (SRA) database with an accession number SRP218063.

## AUTHOR CONTRIBUTIONS

QS designed and supervised the research. LL processed and analyzed all of the data and performed the experiments. LL wrote the article. WJ Y made critical comments and proof read the article. All authors read, edited and approved the final version for publication.

## ACKNOWLEDGEMENTS

This work was supported by the National Science Foundation of China (32201643).

## SUPPORTING INFORMATION

Additional Supporting Information may be found in the online version of this article.

Table S1 The identified flavonoid and their accumulation level in *E. ulmoides* leaves.

Table S2 full-length sequences statistics.

Table S3 Annotation of all expressed genes in *E. ulmoides* leaves.

Table S4 The expression profiles of all identified *E. ulmoides* genes.

Table S5 The evaluation statistics of ATAC sequencing data.

Table S6 Overview of mapping of the ATAC sequencing read

Table S7 The cell preferentially enriched peaks.

Table S8 List of primers used for qRT-PCR analysis.

